# Parallel morphological and functional development in the Xenopus retinotectal system

**DOI:** 10.1101/2025.05.14.654067

**Authors:** Vanessa J. Li, David Foubert, Anne Schohl, Edward S. Ruthazer

## Abstract

The retinotectal projection in *Xenopus laevis* is topographically organized. During the early development of the Xenopus visual system, the optic tectum increases considerably in volume, and retinotectal axons and dendrites undergo extensive activity-dependent remodeling. We have previously observed marked changes in the three-dimensional layout of the tectal retinotopic functional map over the course of a few days. This raised the question of whether such functional reorganization might be attributable to the migration and structural remodeling of tectal neurons as the brain grows. To examine changes in map topography in the context of individual tectal neuron morphology and location, we performed calcium imaging in the optic tecta of GCaMP6s-expressing tadpoles in parallel with structural imaging of tectal cells that were sparsely labelled with Alexa 594-dextran dye. We performed functional and structural imaging of the optic tectum at two developmental time points, recording the morphology of the dextran-labelled cells and quantifying the changes in their positions and the spanning volume of their dendritic fields. Comparing anatomical growth to changes in the functional retinotopic map at these early stages, we found that dendritic arbor growth kept pace with the overall growth of the optic tectum, and that individual neurons continued to receive widespread visual field input, even as the tectal retinotopic map evolved markedly over time.

## Introduction

*Xenopus laevis* has historically been a popular model organism in which to study neurogenesis and topographic wiring of neurons in developing circuits (Cline & Kelly, 2012; Liu et al., 2016; McFarlane & Lom, 2012). In a previous study (Li et al., 2022), we mapped the three-dimensional (3D) layout of the visuotopic map in the tadpole tectum through calcium imaging over developmental stages 42 through 48 (Nieuwkoop & Faber, 1994), a period of robust growth and circuit refinement for the retinotectal system (Gaze et al., 1972; Holt & Harris, 1983; Kutsarova et al., 2017; Sakaguchi & Murphey, 1985). We saw a marked change in the layout of functional retinotopic maps between stages 45 and 48, with the 3D axes of retinotopic organization undergoing large spatial rotations, prompting questions on how the underlying circuit changed during this process.

Starting from the developmental stages when retinal ganglion cell (RGC) axons first reach the tectum until metamorphosis, both the eye and tectum of the tadpole expand and add cells, requiring the visual system to continuously rewire its circuit to maintain visual encoding functionality so that the tadpole can effectively locate food and avoid predators. To add to the complexity of the problem, the patterns of cell proliferation in the retina and tectum do not match: The retina expands radially by adding cells at its ciliary margin (Beach & Jacobson, 1979; Hollyfield, 1971; Straznicky & Gaze, 1971); meanwhile the tectum adds new cells at its caudomedial end, displacing older cells toward the rostral end of the tectum (Herrgen & Akerman, 2016; Lazar, 1973; Straznicky & Gaze, 1972). This means that to maintain an orderly retinotopic map, the circuit must constantly undergo some form of rewiring, by either dynamically shifting physical RGC-to-tectal neuron connections or by functionally changing the receptive field input strengths onto tectal neurons (Chung et al., 1974; Gaze et al., 1979).

In the present study, we sought insights into this rewiring process by following the morphological and functional changes in individual tectal cells and putting them into the context of whole-tectum functional retinotopic map changes. We applied our previously established methods to express the fluorescent calcium indicator GCaMP6s in the tadpole tectum to perform calcium imaging to extract retinotopic maps (Li et al., 2022), but also introduced red fluorescent labelling of individual tectal cells through single-cell electroporation (Haas et al., 2002; Haas et al., 2001). This allowed us to examine developmental changes in the morphology of the labelled cell and its position in the tectum, as well as the functional retinotopy of visual inputs to its dendritic arbor.

## Results

### Characterizing morphological and functional changes in the growing tadpole tectum

We generated tadpoles expressing the genetically-encoded calcium indicator GCaMP6s (Chen et al., 2013) through mRNA blastomere injection, as described previously (Li et al., 2022). This allowed us to visualize the growing tadpole tectum and analyze its functional properties by recording responses to visual stimuli and extracting visuotopic maps. Injecting GCaMP6s mRNA into one blastomere of two-cell stage embryos produced hemi-mosaic animals with GCaMP expression restricted to one lateral half of the animal. In Xenopus tadpoles, RGC axons from one eye project almost exclusively to the contralateral tectum (Munz et al., 2014). Therefore, the tectal hemisphere in the lateral half with GCaMP expression will contain GCaMP in tectal cells but not incoming RGC axons, ensuring the calcium signal recorded from this tectal lobe primarily reflects postsynaptic neuronal activity. When the animals reached stage 44, we performed single-cell electroporation to fill a single or small number of tectal neurons with Alexa Fluor 594 dextran dye. This allowed us to observe and follow the morphology of individual tectal cells and their relative anatomical positions in the tectum over time. We performed two-photon calcium imaging on tadpoles at stage 45-46, then again at stage 48 to compare developmental differences in the same animals (Fig. 1a, c). During each imaging session, we collected high-resolution z-stack images of the tectum to capture the morphology of both the tectum and Alexa 594 dextran-labelled cell (using basal GCaMP6s fluorescence to visualize overall tectal morphology), then performed retinotopic mapping by presenting drifting visual stimuli and recording the evoked GCaMP signal as previously described (Li et al., 2022).

**Fig 1:**
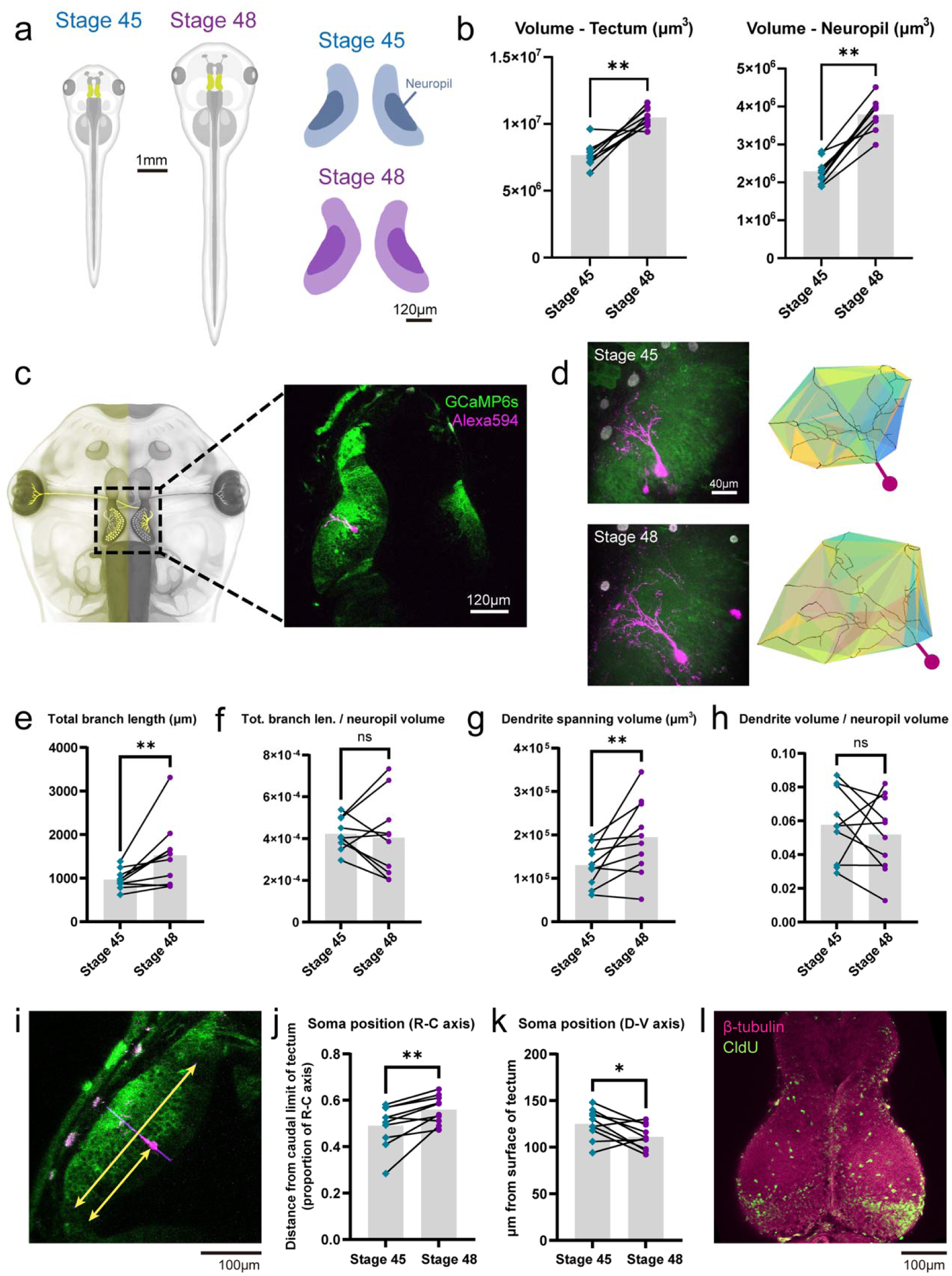
Morphometric changes in the tadpole tectum between stages 45 and 48. **(a)** Illustration of the tadpole and tadpole tectum at stages 45 and 48. The neuropil of the tectum is represented by darker colors. **(b)** Volume of the tectum and tectal neuropil increased significantly from stage 45 to 48. **(c)** (Left) Schematic of a tadpole with hemimosaic GCaMP6s expression restricted to the left half of the animal. (Right) Two-photon image showing GCaMP6s expression limited to tectal cells in the left and RGC axons in the right tectal hemisphere. A single postsynaptic cell in the left tectal hemisphere was labelled with Alexa Fluor 594 dextran fluorescent dye. **(d)** (Left) Maximum projection images of the left tectal hemisphere of the same tadpole at stages 45 and 48, with a single postsynaptic tectal neuron labelled with Alexa 594 dextran. (Right) Reconstructions of the dendritic tree of the dextran-labelled cell, with colored patches showing the boundaries of its 3D spanning volume. **(e)** Dendritic total branch length increased from stage 45 to 48. **(f)** Dendrite density, calculated as total branch length divided by neuropil volume, did not change significantly. **(g)** Dendrite spanning volume increased from stage 45 to 48. **(h)** Dendrite coverage, calculated as dendrite spanning volume divided by neuropil volume, did not change significantly. **(i)** Position of the cell soma along the rostrocaudal (R-C) axis was calculated as the distance from the soma to the caudal limit of the tectum divided by the full length of the R-C axis. **(j)** Labeled neuronal somata shifted rostrally from stage 45 to stage 48. **(k)** Position of the cell soma along the dorsoventral (D-V) axis. From stage 45 to stage 48, cell somata shifted deeper below the dorsal surface of the tectum. **(l)** β-tubulin and CldU labelling in the tectum of an animal treated with 10 mM CldU for 2 h at stage 45 and sacrificed at stage 48. CldU labelling can be seen distributed in the cell body layer in a laminar fashion, revealing the linear proliferation pattern of tectal neurons. CldU puncta can also be seen sparsely distributed in the tectal neuropil. All paired comparisons were done using Wilcoxon matched pairs test, n=10 animals, *p < 0.05, **p < 0.01

### Morphometric changes in the tectum and postsynaptic tectal cell dendritic fields

We manually segmented the tectum and tectal neuropil to acquire volumetric measurements (Fig. 1b), and generated automated 3D reconstructions of the dendritic trees of dextran-labelled tectal neurons to calculate the total branch length and spanning volume of dendritic fields (Fig. 1d).

Between stage 45 and stage 48, a period of extensive neurogenesis (Gaze et al., 1972; Herrgen & Akerman, 2016), there was a significant increase in the volume of the tectum and tectal neuropil (Fig. 1b). Total branch length and spanning volume of the dendritic fields of individual tectal neurons increased significantly (Fig. 1e, g), and Sholl analysis showed an increase in branch complexity (Supplementary Fig. S1a). However, we did not detect a significant change in dendritic branch length per total neuropil volume nor in dendrite coverage (dendrite spanning volume divided by total neuropil volume), indicating that there was not a significant change in the proportion of total neuropil covered by a single cell’s dendritic field due to matched growth of the dendritic arbor and tectal volume (Fig. 1f, h).

We measured the soma position of the dextran-labelled cells along the long axis of the tectal lobe, and found that from stage 45 to 48 the cell somata consistently shifted rostrally and ventrally (Fig. 1i-k), consistent with new tectal cells being added from the caudal end of the tectum (Herrgen & Akerman, 2016). To confirm the pattern of proliferation of tectal neurons, we performed chlorodeoxyuridine (CldU) labelling to label cells that were dividing at the time of administration (Fig. 1l). CldU administered for 2 hours at stage 45 resulted in labelling distributed in a laminar fashion in the middle of the cell body layer of the tectum at stage 48, consistent with tectal cells being added linearly in the tectum.

Previous studies of Golgi impregnated cells in slightly older animals reported that morphological features of tectal cells may be correlated to the position of the cell along the rostrocaudal (R-C) axis due to more mature cells being situated in more rostral positions (Lazar, 1973; Wu et al., 1996). However, for the cells in our dataset we observed no consistent correlation between soma position in the tectum and morphological complexity, including dendrite spanning volume, total branch length, dendrite coverage or dendrite growth rate (Supplementary Fig. S1b-d).

### Changes in functional retinotopic representations in the tectal neuropil

To characterize the functional representation of the visual field in the tectum, we presented tadpoles with repeated drifting bar stimuli across the azimuth and elevation axes of the visual field and recorded calcium responses (ΔF/F_0_) in the tectum to estimate receptive field (RF) positions. A Fourier power spectrum analysis of the calcium signal fluctuations at the stimulus repetition frequency (0.05 Hz) was used to extract the stimulus-driven neural responses. The phase of the response signal at the stimulation frequency corresponds to the time of the peak response and thus the receptive field position on the stimulation screen. Retinotopic mapping was performed by rapidly imaging at multiple depths in one tectal hemisphere to obtain a 3D functional visuotopic map. We regularly observed marked changes in the 3D layouts of azimuth and elevation topographic gradients in the same animal between stage 45 and stage 48 (Fig. 2a-c).

**Fig 2:**
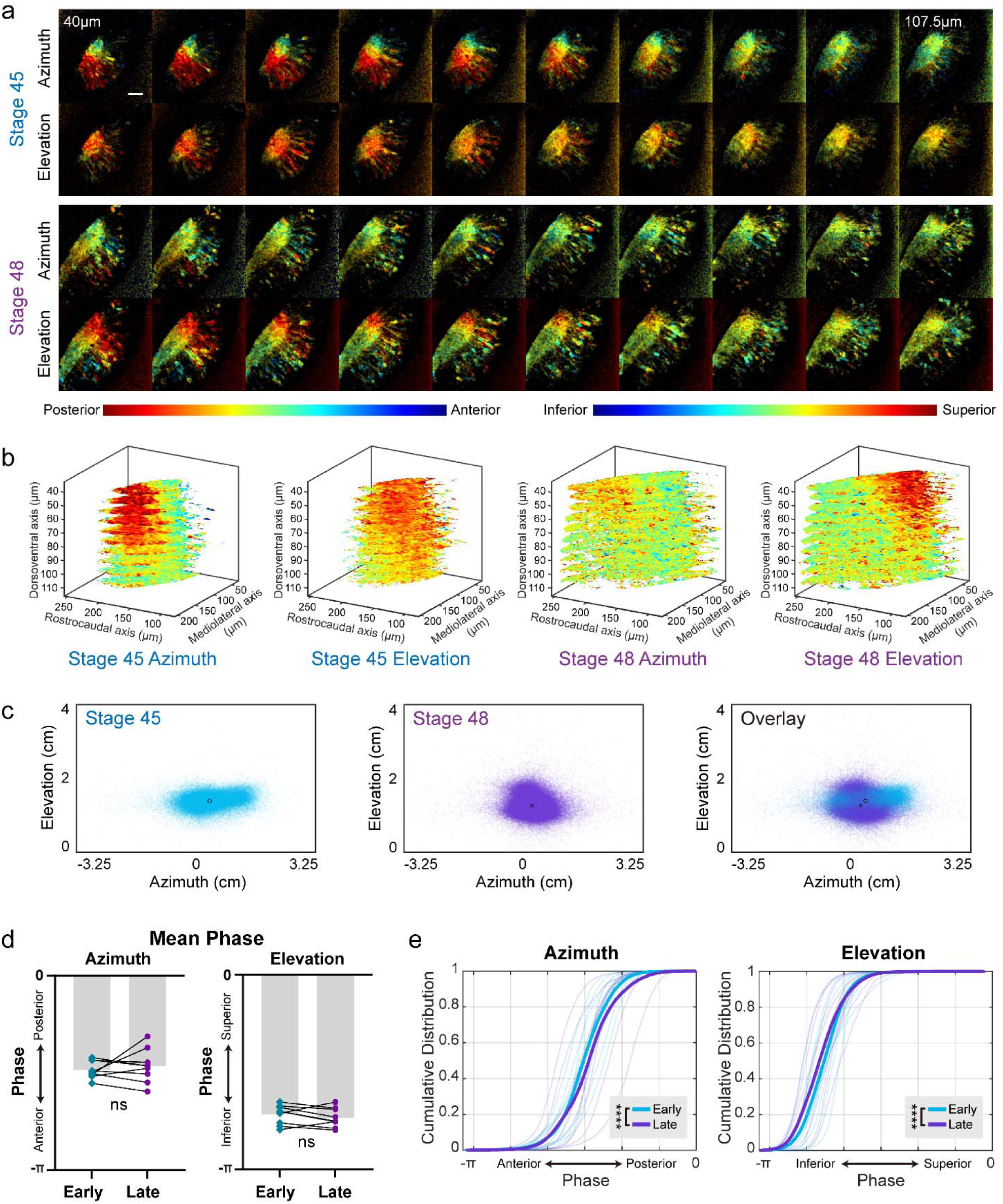
Visual receptive fields represented in the tectal neuropil at different developmental stages. **(a)** Azimuth and elevation receptive field maps from the same animal at stage 45 and stage 48. Receptive field (RF) positions were calculated as the phase of the response to a repeated drifting bar stimulus in the corresponding axis. Pixel intensities indicate signal-to-noise ratio (SNR). Images were taken in stacks of 10 optical sections starting at approximately 40 μm depth from the surface of the tectum, with 7.5 μm between sections. Scale bar is 40 μm. **(b)** 3D renderings of the phase maps from (a), showing only voxels in the neuropil with SNR > 2. **(c)** Neuropil receptive field positions from (b) (all optical sections) mapped onto the stimulus display field. **(d)** Mean neuropil phase did not differ between early and late stages for both azimuth and elevation axes (Wilcoxon matched pairs test, n=9 animals) **(e)** Cumulative probability distribution of neuropil receptive field phase values. Thin lines show data from individual animals (n=9), down-sampled to 2000 random datapoints for each animal. Thick lines show pooled data from all animals. Pooled data show a small but significant shift in the RF distributions between early and late stages for both azimuth and elevation (Kolmogorov-Smirnov test, ****p < 0.0001).

Using the dextran-labelled cells as fiducial markers in the tectum, we matched 3 optical sections containing the labelled cells at early (stage 45-46) and late (stage 48) stages, with 15 μm depth between optical sections, and quantified the topographic gradients along the R-C axis of the neuropil for each optical section (Supplementary Fig. S2a-c). The dorsalmost sections were 40∼70 μm from the dorsal surface of the tectum, depending on the position of the labelled cell. The topographic gradients, as measured by the slope of the linear regression fitted to the RF values, differed considerably between early and late stages. In the dorsalmost section imaged, the gradient of the elevation axis map consistently shifted toward alignment with the R-C axis from early to late stages. However, in deeper optical sections, the directions that topographic gradients shifted were not consistent between animals (Supplementary Fig. S2d). This observation indicates a high degree of lability of the retinotopic map during these developmental stages.

Despite the striking lability in the layout of the topographic gradients within the tectum between early and late stages, when we compared total visual field representations throughout the tectal neuropil for each animal, there was no overall shift in the mean receptive field center of either the azimuth or elevation axes (Fig. 2d). However, the overall distribution for all measured RFs showed a slight but significant shift towards higher phase values in azimuth (corresponding to more representations in the posterior field) and lower phase values in elevation (corresponding to more representations in the inferior field) in late-stage recordings (Fig. 2e).

Given the surprising amount of map reorganization observed, we wondered how the inputs to individual tectal neurons might be changing across these time points. Therefore, we next examined the RFs of voxels colocalizing with the dendritic arbors of the dextran-labelled postsynaptic cells (Fig. 3a) and their relationship to the RF coverage in the rest of the tectal neuropil. Cumulative distribution plots of pooled RF phase values from voxels colocalized with the dendritic fields of dextran-labelled cells showed a developmental shift towards more coverage in anterior visual fields (lower phase values) in azimuth and a small shift towards superior visual fields (higher phase values) in elevation (Fig. 3b) at the later stage, opposite to the trend that was seen for the whole neuropil (Fig. 2e). This points to a scenario where the RFs of individual tectal cells are shifting gradually, but this change in mature cells may be offset by newer cells being added to the tectum, preserving the overall RF distribution in the whole tectum.

**Fig 3:**
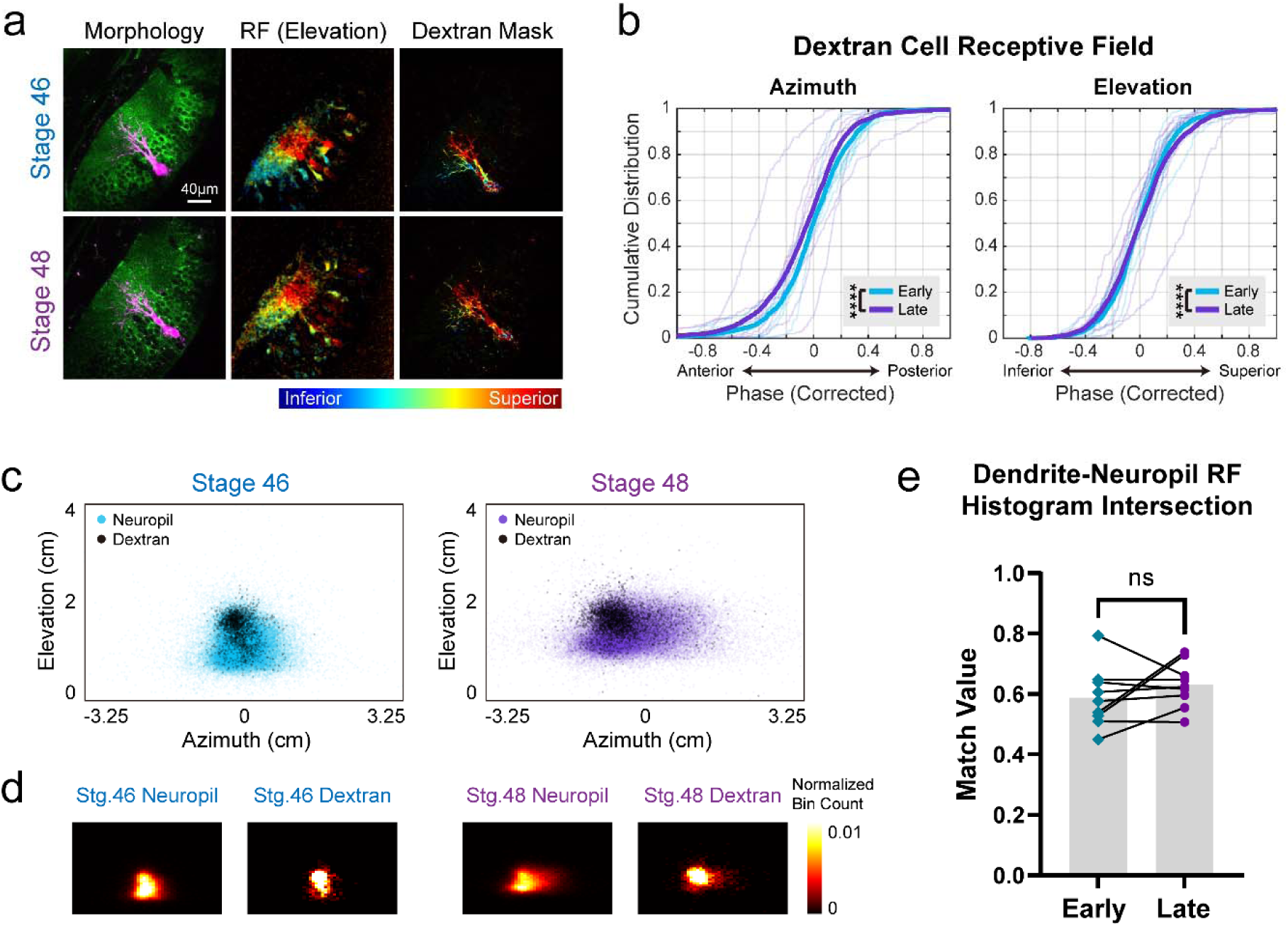
Receptive fields measured at the dendrites of single tectal neurons at different developmental stages. **(a)** Example optical section from the same animal imaged at stage 46 and 48: (Left) Morphology of the tectum with a single-cell electroporated dextran-labelled neuron. (Middle) Elevation receptive field maps (pixel intensities indicate SNR). (Right) Elevation receptive field maps overlaid with mask of dextran labelling. **(b)** Cumulative distribution of receptive field phase values recorded from areas in the neuropil with dextran labelling (data from 10 optical sections for each animal). Thin lines show data from individual animals (n=9), down sampled to 200 datapoints for each animal. Thick lines show pooled data from all animals. Phase values were corrected by mean centering (see Methods). Pooled data from all 9 animals show significant difference between early and late stages for both azimuth and elevation (Kolmogorov-Smirnov test, ****p < 0.0001). **(c)** RF phase values from the animal in (a) (data from 10 optical sections) mapped onto the stimulus display field. Colored scatter points represent data points from the neuropil; black scatter points represent datapoints from areas in the neuropil with dextran labelling. **(d)** Heat maps showing the data from (c) binned into 2D histograms (50 x 50 bins. Bin counts were normalized so that the sum of all bin counts in each histogram equals 1). **(e)** The match value (see Methods) between RF histograms for neuropil and dextran-labelled cells was high, and didn’t change significantly between early and late stages (Wilcoxon matched pairs test, n=9 animals).

The fact that we did not see a morphometric change in the dendritic coverage of labelled cells as a proportion of total neuropil volume (Fig. 1h) led us to ask if this would also be reflected functionally in RF representations in the dextran-labelled cells compared to the whole neuropil. We binned RF phase values recorded from the total neuropil and dendrites of dextran-labelled cells into 2D histograms, and found a high degree of similarity between the two histograms (Fig. 3c-e). This degree of similarity did not change significantly between early and late stages, suggesting high levels of visual input convergence to each tectal neuron that was preserved over these developmental stages.

### Changes in visual response properties in individual tectal neurons

Receptive field analysis based on drifting bar responses in individual tectal cell soma regions-of-interest (ROIs) yielded retinotopic maps with topographic gradients that were generally consistent with the corresponding neuropil retinotopic maps from the same animals (Fig. 4a). The distribution of cell body RF positions along the neuropil R-C axis largely matched the neuropil RF distribution (Supplementary Fig. S3-4). As in the neuropil, the mean RF phase of all cell body ROIs from the whole tectum of individual animals revealed no significant shift between early and late stages (Fig. 4b), but the overall distribution of RF centers of individual cells showed a very slight but significant posterior shift in azimuth and a small downward shift in elevation (Fig. 4c).

**Fig 4:**
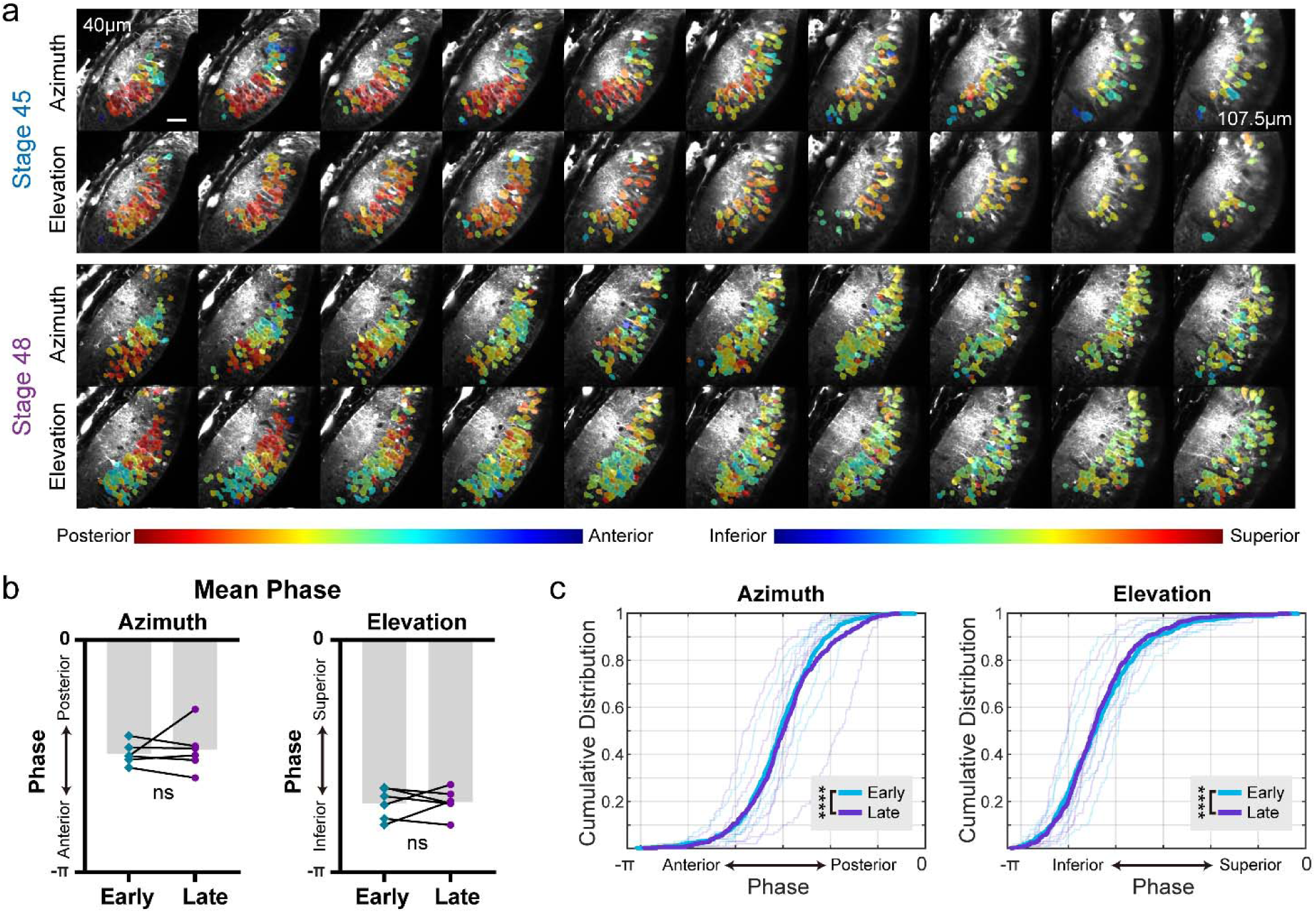
Receptive fields of tectal neuron cell bodies across developmental stages. **(a)** Azimuth and elevation cell body receptive field maps from the same animal and showing the same optical sections as in Fig. 2a. RF phase values were calculated from the mean ΔF/F trace for each cell body ROI. Only cells with SNR > 2 are shown. Scale bar is 40 μm. **(b)** Mean RF phase for each animal (n = 6). No significant difference between early and late-stage animals (Wilcoxon matched pairs test) **(c)** Cumulative probability distributions of cell body RF phase values. Thin lines show data from individual animals (n=6), down sampled to 100 cells for each animal. Thick lines show pooled data from all animals. Pooled data reveal a small difference between early and late stages for both azimuth and elevation (Kolmogorov-Smirnov test, ****p < 0.0001).

Because we expressed GCaMP6s by mRNA microinjection into blastomeres at the 2-cell stage, one possible alternative explanation for the lability of RF gradients and distributions could be that degradation of mRNA and protein over time might lead to less reliable responses in older animals. We therefore quantified the signal-to-noise ratio (SNR) and direction selectivity index (DSI) of cell body responses to drifting bars and their degree of direction preference for drifting bars presented in opposite directions (Fig. 5). Comparing pools of cell body responses recorded at early vs late stages, we saw an increase in SNR at the later stage compared to younger animals. Additionally, we found that direction selectivity significantly decreased with age. Thus, GCaMP6s protein expression was likely sufficiently high at both the early and late experimental time points to obtain robust visual responses, and adequate to reveal developmental changes in stimulus selectivity.

**Fig 5:**
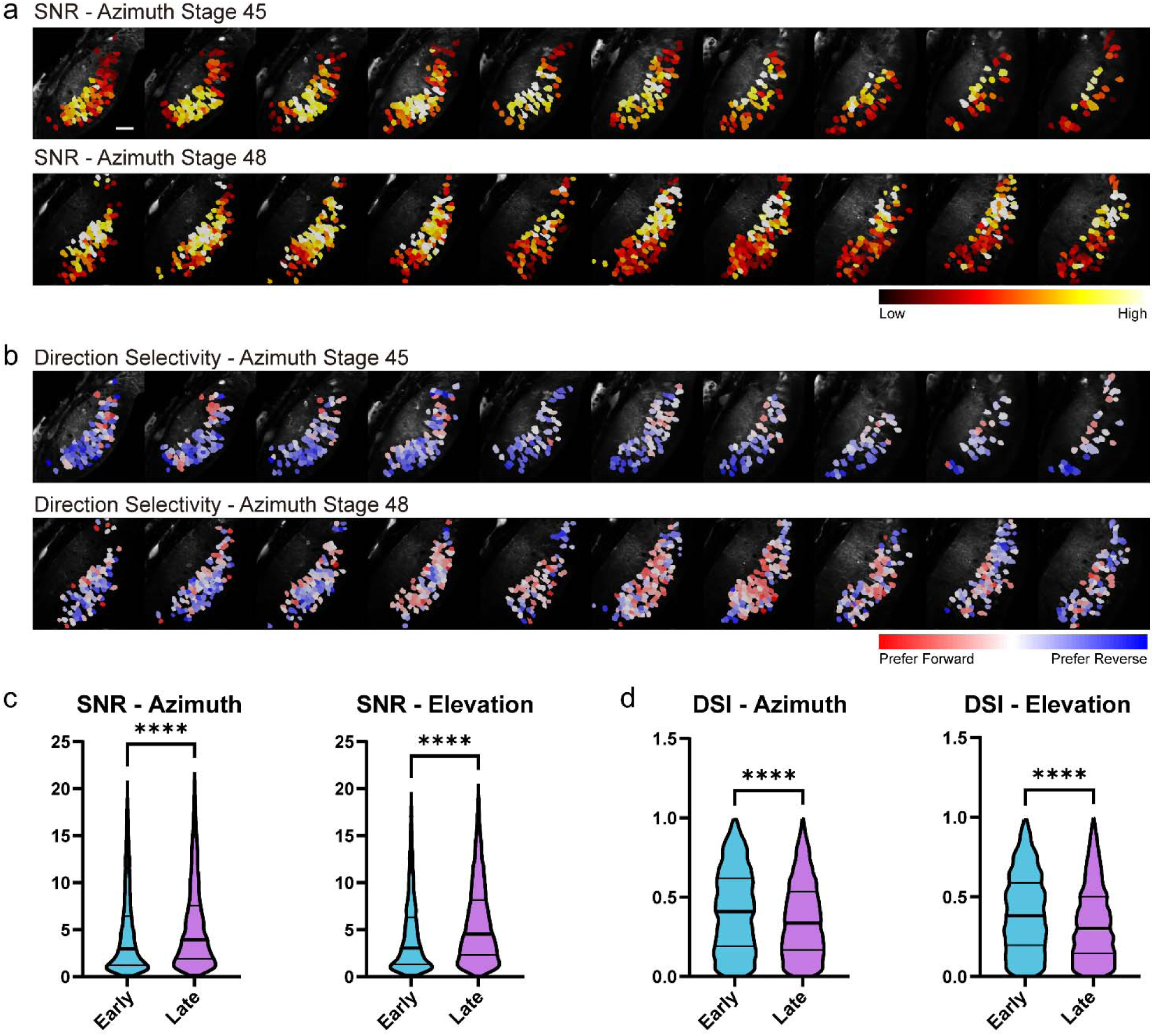
Signal to noise ratio and direction selectivity in postsynaptic tectal cells. **(a)** SNR of cell body responses to drifting bars in the same animal at stage 45 and 48. Scale bar is 40 μm. **(b)** Direction selectivity of tectal cells. Darker colors indicate a higher direction selectivity index (DSI). **(c)** SNR of cell body responses to drifting bars at early (stage 45-46) and late (stage 48) stages, cells pooled from 5 animals. SNR was significantly higher at late stage for both azimuth and elevation (azimuth: n_early_ = 3679, n_late_ = 3781; elevation: n_early_ = 3633, n_late_ = 3886) **(d)** DSI of tectal cells at early and late stages, pooled from 5 animals. Only cells with SNR > 2 were included. DSI was significantly lower at late stage for both azimuth and elevation (azimuth: n_early_ = 2185, n_late_ = 2704; elevation: n_early_ = 2206, n_late_ = 2999). All paired comparisons were performed using two-tailed Mann-Whitney tests, ****p < 0.001.

## Discussion

In this study, we examined the *Xenopus laevis* retinotectal system over a key period of rapid growth in early visual system development. We characterized morphological and functional changes at both the whole-tectum and single-cell levels to gain a better understanding of the potential ways morphology translates to function, and how changes at the single-cell level contribute to changes in the whole circuit. Our method of expressing GCaMP in tadpoles through mRNA blastomere injection produced hemi-mosaic animals with GCaMP expression in one of the two tectal hemispheres restricted to postsynaptic tectal cells, with no GCaMP in the presynaptic retinal ganglion cell arbors. This ensured that the calcium signal recorded from this tectal hemisphere was comprised exclusively of postsynaptic somatic and dendritic components of the retinotectal map, without contamination from presynaptic RGC axonal signals that would instead reflect axonal inputs. Next, by labelling individual neurons in this tectal lobe by single-cell electroporation of dextran-conjugated dye, we established a system where the morphology and functional activity of the whole tectum and individual postsynaptic cells within the tectum could be observed in parallel. We examined the developmental changes in the position, morphology, and functional visual field representations in the labelled individual cells, and compared them to changes seen in the whole tectum.

Our morphometric measurements showed that while both the whole tectum and dextran-labelled postsynaptic dendritic arbors saw significant increases in volume, the proportion of total tectal volume occupied by the dendritic field of the labelled cell remained relatively constant. These observations were echoed in functional aspects: While there were considerable shifts in the layout of retinotopic gradients within the whole tectum, there was relatively little change in the distribution of total visual field inputs to the dendrites of individual labelled tectal neurons.

Notably, regarding developmental refinement in the retinotectal circuit, Sakaguchi and Murphey (1985) reported that while RGC axons continue to grow and increase in complexity over development, their expansion is slower than that of the overall tectal neuropil, thereby resulting in each axon occupying a progressively smaller fraction of the total neuropil over time. In contrast, our morphometric observations of tectal cell dendritic arborization showed ongoing but matched growth of tectal cell dendritic arbors and whole tectal neuropil. Functionally, we found the range of visual field representations found in tectal cell dendritic arbors to be quite wide, and evidence of topographic refinement within the observed developmental stages was subtle. This suggests an overall scenario where the presynaptic portion of the retinotectal circuit is becoming more precise, but the postsynaptic portion maintains a relatively large input range and does not follow the same pattern of refinement as the presynaptic component. Although dendritic arbors of individual tectal cells grow continuously during this period, they cover the same proportion of the tectum and receive retinal inputs that represent the same proportion of the visual field (Fig. 6).

**Fig 6:**
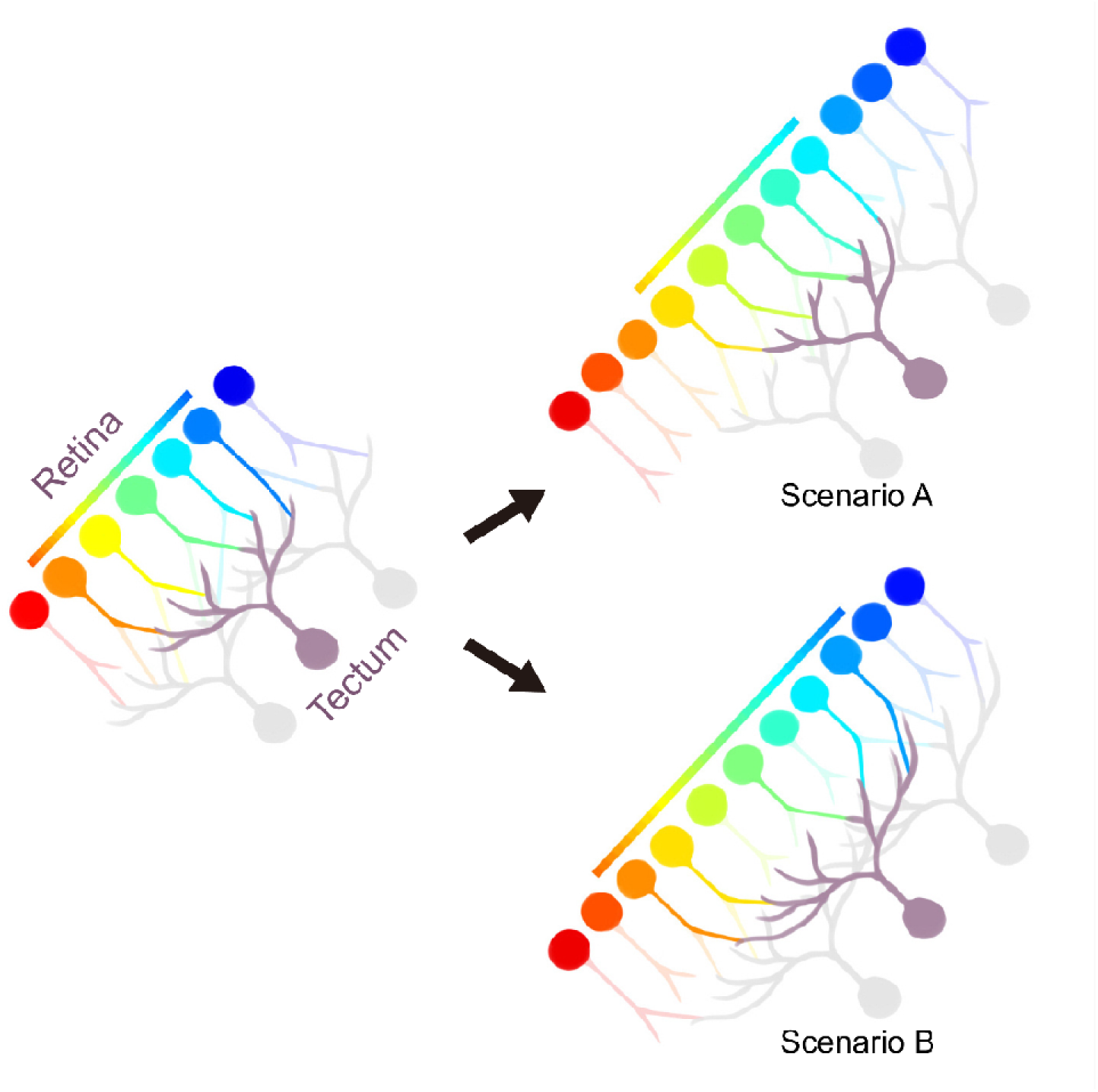
Two possible scenarios for change in retinal inputs to tectal neurons during development. Scenario A: As the retina grows, individual tectal cells receive input from the same number of RGCs, but the RGCs represent a smaller portion of the visual field, leading to reduced input range to the tectal cell. Scenario B: Receptive fields of all RGCs providing input to an individual tectal cell are distributed across the same proportion of the visual field as earlier, leaving the range of visual inputs to each tectal cell unchanged. Our observations of tectal cell dendrites expanding to covering a conserved proportion of the neuropil and displaying a similar range of visual receptive fields between early and late stages point to Scenario B being more likely.

A recent study in adult mouse superior colliculus (Molotkov et al., 2023) found RGC axon terminals to form near-perfect retinotopic tiling in the SC, with retinotopic precision measured in RGC axon terminals found to be higher than that in local SC neuron somata. Based on these results, the authors suggested that retinotopic precision in the SC arises from topographically precise input from presynaptic cells rather than from computational reconstruction performed by postsynaptic circuitry. However, our observations in Xenopus tadpoles suggest that inputs received by tectal cell dendrites in the developing tectum are broad and imprecise, and the animal would likely need to rely on computations in postsynaptic tectal cells to recover sufficient (albeit coarse) visual information to guide behaviour.

The Xenopus tectum in early development faces two prominent challenges that could determine its strategy for encoding and processing visual information. On one hand, computational power in the developing tectum is much more limited, both in terms of the total number of available tectal neurons and the ability of these neurons to resolve spatial information. In the developmental stages examined in this study, the volume of the tectum is only about one-hundredth of what it will grow to in adult animals (Gaze et al., 1974; Roth & Walkowiak, 2015), and tectal neurons at these stages have been shown to have very large receptive fields (Dong et al., 2009; Gaze et al., 1974; Hiramoto & Cline, 2024; Van Horn et al., 2017; Vislay-Meltzer et al., 2006). On the other hand, the developing tectum needs to reconcile with the constant integration of newly proliferated cells into the tectal circuit and the resulting need to adjust local connections to maintain stable and behaviourally relevant visual processing output. With individual tectal neurons lacking the ability to encode precise retinotopic information through single stimulus-evoked responses, the animal likely needs to integrate the responses of multiple tectal cells to recover spatial relationships. Under these circumstances, the topographically precise wiring of retinal inputs to tectal targets become less important, which could explain the apparent lack of topographic refinement observed in tectal cell dendrites over a short period of time. An inherently wide input range and reduced requirement for topographically precise connections can in turn become beneficial when integrating new cells into the circuit: the amount of branch rewiring necessary to reestablish working order in the circuit would be reduced, cutting down on metabolic expenses involved in the process.

It should be noted that in the present study, our measurements of calcium signals from voxels overlapping with dendrites of dextran-labelled tectal cells are likely a close reflection of the range of retinal inputs available to that cell, but they do not give us precise information on the computational output from that tectal cell, which would be most reliably measured at the cell soma. Functional analyses on somatic signal from the dextran-labelled cells in our dataset was unfortunately not possible as a large proportion of the dextran-labeled tectal neurons had poor GCaMP signal strength from their cell bodies. We performed our single-cell electroporations blind to GCaMP responsiveness, resulting in many dextran-labeled cells that had little or no somatic calcium signal. The harsh process of membrane disruption required for electroporation may have also perturbed GCaMP activity within the cell. However, based on the increased SNR we measured from somatic responses of other cells in the tectum, we expect that had we been able to evaluate somatic responses in these dextran-labelled cells, we would see an increase in their ability to reliably decode visual information despite the lack of topographic refinement in the input they are receiving. We also expect that later in development, as the size of the tectal neuron population surpasses a certain threshold, the system would switch from favoring broad integration over diffuse inputs to favoring focused connections from selected inputs, at which point developmental refinement of topographic precision in tectal neurons would become much more prominent.

## Materials and Methods

### Animals

All procedures were approved by the Animal Care Committee of the Montreal Neurological Institute at McGill University and carried out in accordance with Canadian Council on Animal Care guidelines.

Female albino *Xenopus laevis* frogs (RRID: XEP_Xla300) from our in-house breeding colony were injected with pregnant mare serum gonadotropin (Prospec) and human chorionic gonadotropin (Sigma-Aldrich) to induce ovulation. Eggs were collected and fertilized with sperm from male albino X. laevis frogs. Blastomere microinjection of GCaMP6s mRNA to create animals with hemilateral GCaMP expression was performed as previously described (Li et al., 2022). Briefly, a solution of purified GCaMP6s (500 pg) mRNA in 2 nl RNase-free water was pressure injected into one blastomere of two cell–stage embryos using a calibrated glass micropipette attached to a PLI-100 picoinjector (Harvard Apparatus). GCaMP mRNA was prepared by cloning the coding sequence of GCaMP6s into pCS2+, then linearizing plasmids with Notl, and transcribing the capped mRNA of GCaMP6s with the SP6 mMessage mMachine Kit (Ambion; Thermo Fisher). Developing animals were screened to select individuals with bright and unilaterally restricted GCaMP fluorescence. When these animals reached stage 44, single-cell electroporation was performed on the tectal hemisphere with postsynaptic GCaMP expression to label a single or small group (2∼3) of postsynaptic tectal cells with Alexa Fluor 594 dextran.

All tadpoles were reared in biological oxygen demand incubators on a 12/12-h light/dark cycle. Rearing medium was 0.1x Modified Barth’s Solution with 4-(2-hydroxyethyl)-1-piperazineethanesulfonic acid (HEPES) buffer. Tadpole developmental stages were determined according to Nieuwkoop and Faber (1994). Xenopus tadpoles do not exhibit sexually dimorphic traits at the stages used in this study.

### CldU labelling and whole mount immunofluorescence

CldU labelling and immunofluorescence was performed based on McKeown and Cline (2019). Animals were placed in rearing solution with 10mM CldU (Sigma-Aldrich) for 2 h at stage 45, then sacrificed at stage 48. Brains were extracted and fixed with 4% paraformaldehyde in phosphate-buffered saline (PBS, Sigma) overnight at 4°C, washed in PBS, then incubated in PBS-T (0.1% Triton X-100 (VWR) in PBS) for 10 min, followed by 0.5% Triton X-100 in PBS for 1 h. Samples were then treated with 2N hydrochloric acid at 37°C for 20 min for epitope retrieval, rinsed for 20 min 3 times in PBS-T, then placed in blocking buffer (1% bovine serum albumin (Fisher) and 5% normal goat serum (Sigma) in PBS-T) for overnight at 4°C. Next, the samples were incubated with primary antibodies (rat anti-BrdU, Abcam, 1:500) for CldU, and β-tubulin (chicken anti-β-tubulin, Chemicon, 1:1000) diluted in blocking buffer for 3 days at 4°C, washed in PBS-T for 20 min 3 times, then incubated in secondary antibodies (Alexa Fluor 555 goat anti-rat (Invitrogen) for BrdU, Alexa Fluor 488 goat anti-chicken (Invitrogen) for β-tubulin) overnight at 4°C. Finally, the brains were washed with PBS-T, mounted with half a drop of Aqua-Poly/Mount (Polyscience) onto slides fitted with custom mounting spacers (4 layers of adhesive tape (Scotch 3M, 810R-1833) with a hole punched in the center of the tape square) and sealed with a coverslip for imaging.

### In vivo imaging

Fluorescence images were captured with a high-speed resonance scanner-based two-photon microscope (Thorlabs) with piezoelectric focusing (Physik Instrumente) on a 1.0 NA 20x water immersion objective (Nikon). Emission signal was collected in 2 channels through 525/50 and 630/92 bandpass filters, respectively. For morphometric imaging, an excitation wavelength of 840 nm was used to simultaneously image GCaMP6s and Alexa Fluor 594. For functional imaging with GCaMP6s, an excitation wavelength of 910 nm was used. For imaging CldU immunofluorescence, 990 nm was used for Alexa Fluor 488 and 555.

### Morphometric imaging and analysis

Tadpoles were imaged at stage 45 and again at stage 48. In each imaging session, tadpoles were anesthetized by immersion in 0.02% MS-222 (Sigma-Aldrich) and placed on a block of Sylgard 184 silicone elastomer (Dow Silicones) with a carved indentation to hold the animal immersed in a small amount of medium. Stacks of 512 x 512 pixel images stepping through the tectum in 2 µm steps were collected at two magnifications: low magnification 1.492 µm per pixel to show both tectal hemispheres; high magnification 0.482 µm per pixel to focus on one tectal hemisphere. Low magnification images were first denoised using the CANDLE algorithm (Coupé et al., 2012) implemented in MATLAB (MathWorks, RRID: SCR_001622), then segmented in 3D Slicer (Fedorov et al., 2012) to obtain volume measurements of the tectum and tectal neuropil. High magnification images were used to reconstruct the dendritic trees of the dextran-labelled cells using the TREES Toolbox (Cuntz et al., 2010) in MATLAB. Total branch length, dendrite spanning volume and Sholl analyses of the reconstructed dendritic trees were calculated using TREES Toolbox functions. For all morphometric analyses, we considered 180 µm depth from the dorsal surface of the tectum to be the boundary between the tectum and tegmentum, and only structures above this boundary were analyzed.

### Functional imaging and visual stimulation

Tadpoles were immobilized by immersion in 2 mM pancuronium dibromide (Sigma) and embedded in 1% low melting point agarose in a custom chamber with a glass coverslip window on one side, through which the animal could view visual stimuli presented on an LCD screen. The LCD display area measured 6.5 cm (width) × 4 cm (height). The tadpole was positioned so the eye was 2.2 cm from the screen, aligned to the center of the bottom edge of the display area. From this viewpoint, the display area spans roughly 110° of visual angle in azimuth and 80° in elevation. A #29 dark red Wratten filter (Kodak) was installed on the LCD screen to prevent light from the display from interfering with the calcium signal. Custom MATLAB scripts based on the Psychophysics Toolbox (Brainard, 1997; Kleiner et al., 2007; Pelli, 1997) (RRID: SCR_002881) were used to generate the visual stimuli and synchronize stimulus presentation with image capture. Visual stimuli were presented monocularly, and calcium signal was imaged from the tectum contralateral to the stimulated eye. Images were captured by rapidly scanning through 10 optical sections separated by 7.5 µm at a rate of 4.5 Hz per section and 256 × 256-pixel resolution (0.963 µm per pixel).

### Receptive field mapping and analysis

Processing and analyses of calcium imaging data were performed with custom scripts in MATLAB and Fiji (RRID: SCR_002285). Image alignment and motion correction were performed with either the MATLAB imregtform() library function, NoRMCorre (Pnevmatikakis & Giovannucci, 2017) implemented in MATLAB, or Suite2P. For neuropil analyses, masks were manually drawn for the neuropil area and only pixels within the masked area were analyzed. For cell body analyses, ROI segmentation and fluorescence trace extraction were performed with Suite2P (Pachitariu et al., 2016).

#### Receptive field mapping

Receptive field mapping using drifting bar stimuli was performed as previously described (Li et al., 2022). Briefly, a mapping stimulus consisting of a single vertical or horizontal black bar drifting at a constant rate along the full span of the anterior-posterior or superior-inferior axis of the LCD screen was presented repeatedly at regular intervals while recording calcium responses in the tectum. A Fourier transform was applied to the first differential of response traces, the Fourier component at the stimulus frequency yielding the amplitude and phase of the peak response to the stimulus. The phase of the peak stimulus response converts to receptive field position.

To correct for latency in the response due to the continuous activation of retinotectal neurons by the stimulus, mapping experiments were conducted in pairs of trials where the stimulus bars sweep in opposite directions, then a difference was taken between the response phases from the two trials to obtain an absolute response phase. For an experiment with an interval of t_blank_ between repeated drifting bar presentations, absolute phase φ^+^ can be found by

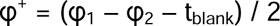

where φ^1^ and φ^2^ are the relative phases in the two opposite directions.

#### Signal to noise ratio

Signal to noise ratio (SNR) of a stimulus-evoked response is defined as A_r_/σ, where A_r_ is the amplitude of the Fourier component at the stimulus frequency, and σ is the standard deviation of all amplitudes at frequencies above the stimulus frequency. When using SNR as thresholds for data analysis and display in Fig. 2 through 5, the smaller of the two SNR values from a pair of mapping experiments was used to exclude noisy voxels that do not show a reliable receptive field representation. For evaluating SNR levels in cell body ROIs in Fig. 5a and 5c, the larger of the two SNR values from a pair of mapping experiments was used to allow us to assess the responsiveness of voxels to visual stimulation.

#### Direction selectivity index

Direction selectivity index (DSI) was calculated for a pair of drifting bar directions (azimuth: forward vs reverse; elevation: upwards vs downwards) as

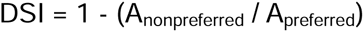

where A_preferred_ is the amplitude of the response to the preferred direction, and A_nonpreferred_ is the amplitude of the response to the opposite direction.

#### Comparing RF phase distributions via histogram intersection

To compare the distribution of RF phase values recorded from the whole neuropil versus the dendrites of dextran-labelled cells, we evaluated the similarity between the two distributions by computing their histogram intersection, as defined by Swain and Ballard (1991).

We mapped RF phase values for all voxels from the neuropil and dextran groups onto the stimulus display field and binned them into 2D histograms with 50 bins along each visual axis (azimuth and elevation), totaling n=2500 bins (Fig. 3c). Bin counts for each individual bin were normalized so that the sum of all bin counts in the 2D histogram equaled 1.

The intersection between the histogram for RF phase values for dextran-labelled voxels, *D*, and the histogram for RF phase values from all neuropil voxels, *M*, is defined to be

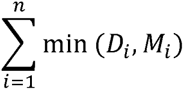

The match value between the two histograms is defined as

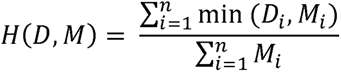

The match value *H(D, M)* is a fractional value between 0 and 1, where higher match values indicate greater similarity between the two distributions.

#### Mean centering correction for phase values

We expected inter-animal variability to affect early-late stage comparisons disproportionately more when looking at the dendritic fields of single cells compared to when looking at the whole neuropil. We therefore performed corrections on the phase data shown in Fig. 3b as follows:

For each animal, calculate the mean phase of all voxels colocalized with the dendritic fields of dextran-labelled cells at early and late stages, X_dex_early_ and X_dex_late_, and the mean phase of all voxels in the neuropil at early and late stages, X_np_early_ and X_np_late_.

For early-stage data, the corrected phase value φ_corrected_ is calculated from the raw phase value φ_raw_ as

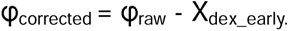

For late-stage data, the corrected phase φ_corrected_ is calculated from the raw phase φ_raw_ as

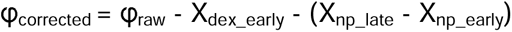

### Statistical analysis

Statistical tests, as indicated in the figure legends, were performed using Graphpad Prism 10 (GraphPad software, RRID: SCR_002798).

## Data and Code Availability

Quantitative data and custom MATLAB code used for analyses and figure generation in this paper can be found at https://figshare.com/projects/Parallel_morphological_and_functional_development_in_the_Xenopus_retinotectal_system/248822 and https://github.com/RuthazerLab/XenMorphFunc.

## Acknowledgements

This work was funded by a Natural Sciences and Engineering Research Council of Canada grant (RGPIN-2024-05306) and Canadian Institutes of Health Research grant (PJT-180478) to ESR, a Molson Neuro-Engineering Award, Shuk-Tak Liang Fellowship and McGill University Integrated Program in Neuroscience Studentship to VJL.

## Supplementary Figures

**Fig S1:**
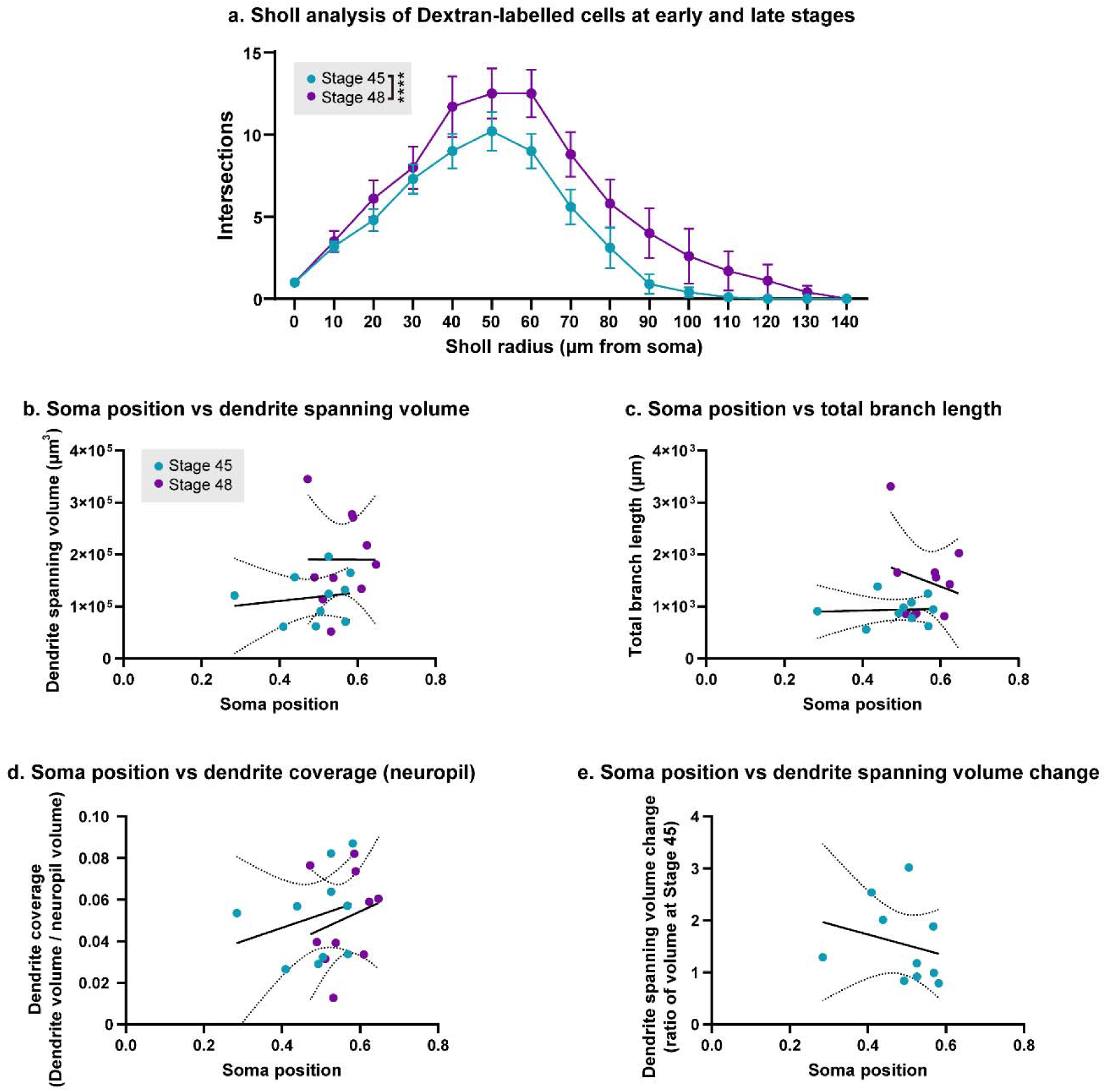
Morphometric changes in dextran-labelled cells between stages 45 and 48. **(a)** Sholl analysis for dextran-labelled cells at Stage 45 and 48 (mean ± SEM number of intersections, n=10 animals). Two-factor repeated measures ANOVA for stage and Sholl radius found significant main effects for both stage and Sholl radius (Stage F(1,135) = 35.55, ****p < 0.0001); Sholl radius F(14,135) = 22.45, ****p < 0.0001). **(b∼e)** Relationships between various morphometric measurements of dextran-labelled cells and the position of their cell soma along the rostro-caudal axis of the tectum. Solid and dotted lines show linear regressions (performed separately for stage 45 and 48 data) and their 95% confidence intervals. **(b)** Soma position vs dendrite spanning volume. Slopes for linear regressions were not significantly different from zero (p_45_ = 0.6594, p_48_ = 0.9950). **(c)** Soma position vs total branch length. Slopes for linear regressions were not significantly different from zero (p_45_ = 0.8643, p_48_ = 0.5366). **(d)** Soma position vs dendrite coverage. Slopes for linear regressions were not significantly different from zero (p_45_ = 0.4535, p_48_ = 0.5310). **(e)** Soma position at Stage 45 vs change in dendrite coverage between Stage 45-48. Slope for linear regression was not significantly different from zero (p = 0.5045).

**Fig S2:**
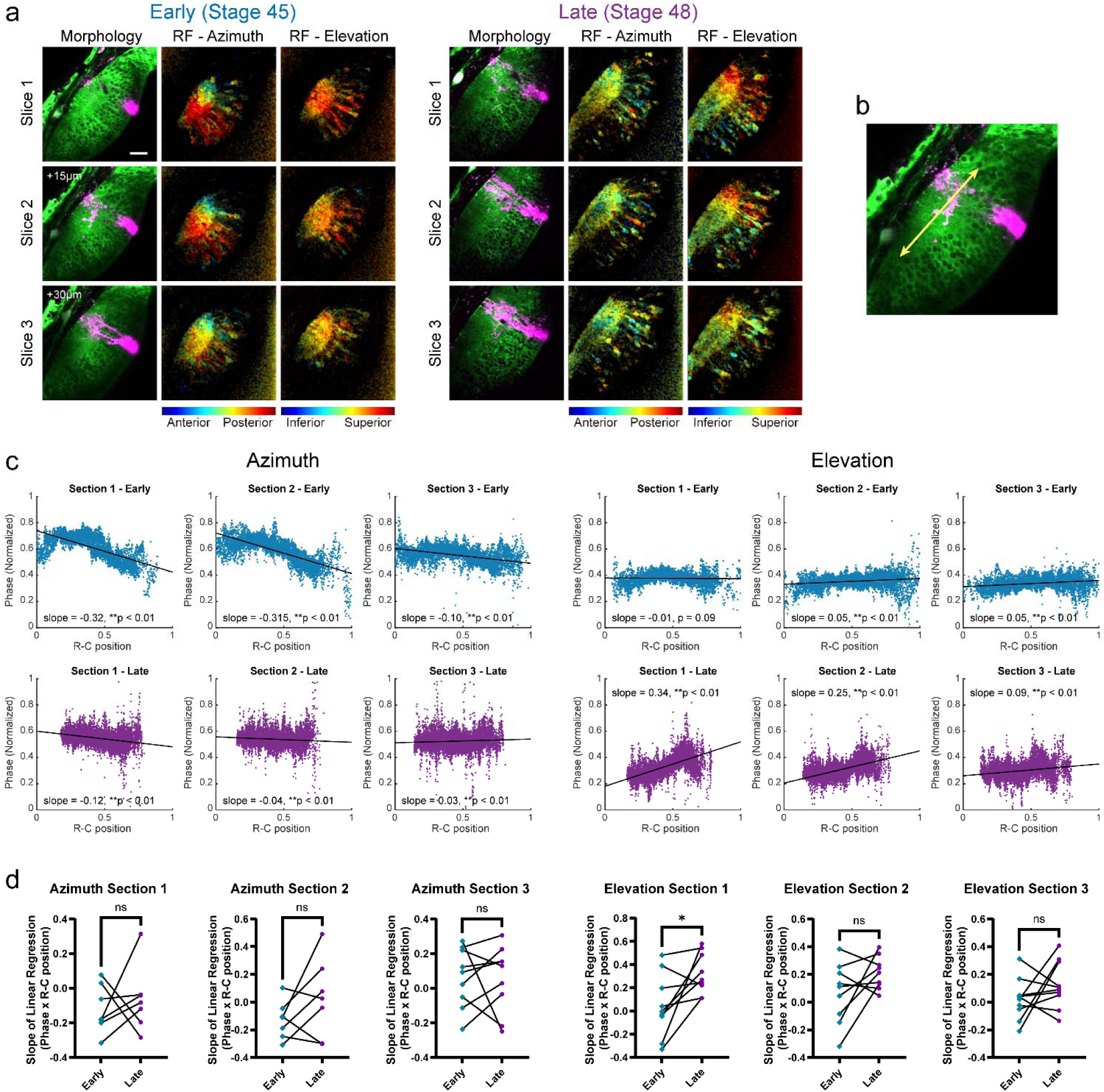
Comparing retinotopic gradients in three matched optical sections from the same animal at early and late stages. **(a)** Example images from one animal. Scale bar in upper left image is 40 μm. The three optical sections are spaced 15 μm in depth from each other, with Section 3 at the bottommost depth. In each row of 3 images for each optical section: (Left) Morphology of the tectum showing dextran-labelled cell. (Middle) Azimuth receptive field map (pixel intensities indicate SNR). (Right) Elevation receptive field map. **(b)** For each stage, the rostrocaudal axis of the neuropil of the center section, Section 2, was used as the reference R-C axis when quantifying the topographic gradients along the R-C axis. An R-C position of 0 is closest to the caudal edge of the neuropil, 1 is closest to the rostral end. **(c)** Distribution of receptive field phase values along the R-C axis for the example animal and optical sections shown in (a). Black line shows simple linear regressions fitted to the data. Phase values are scaled from the original range of [-pi, 0] to a range of [0, 1], therefore the range of the linear regression slope is [-1,1], and a slope of 1 or -1 indicates a topographic gradient that is perpendicular to the R-C axis. **(d)** Comparing slopes of linear regressions fitted to phase vs R-C position (n=10 animals). Data for animals/sections where the linear regression wasn’t a good fit were excluded. Comparing slope values at early vs late stages yielded no significant direction of change for all cases except for the elevation gradients in Section 1, which showed a significant increase (Wilcoxon matched-pairs test, *p = 0.0273).

**Fig S3:**
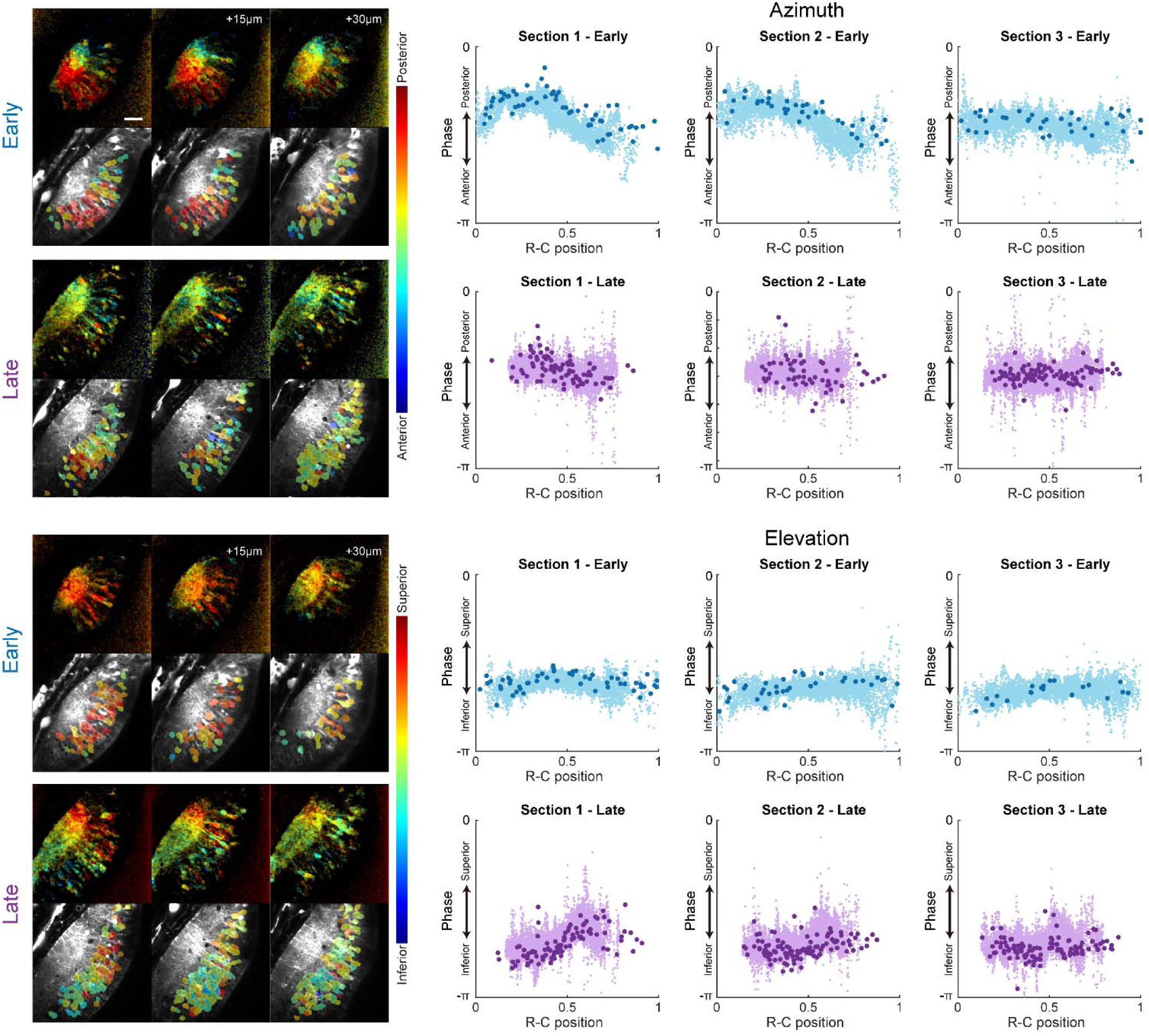
Comparing the distribution of retinotopic representations from the neuropil and cell bodies. Data from same animal and optical sections shown in Fig. S2(a-c). Scale bar in upper left image is 40 μm. Dark points in scatterplots denote data from cell bodies, lighter points denote data from the neuropil.

**Fig S4:**
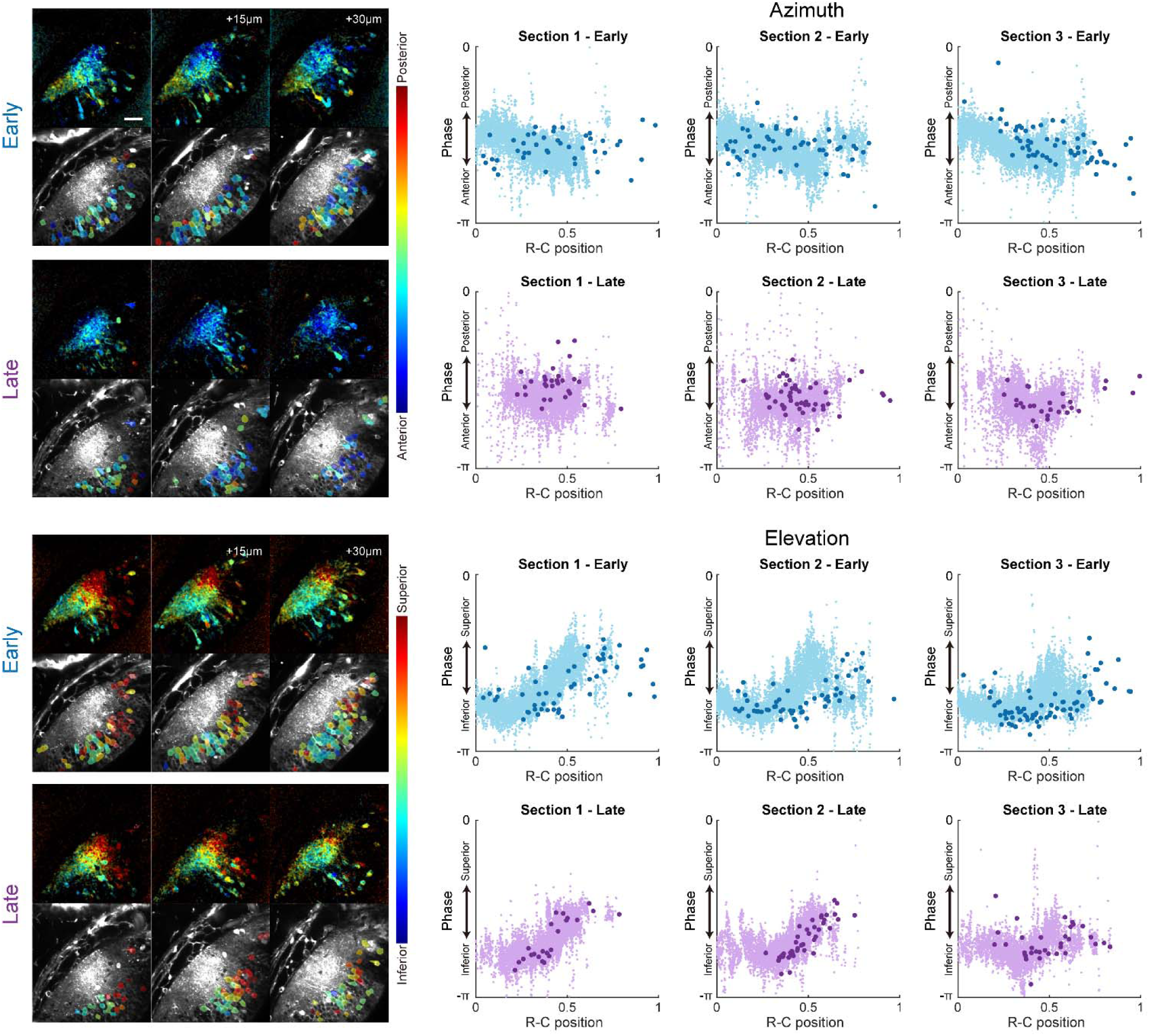
Comparing the distribution of retinotopic representations from the neuropil and cell bodies, showing data from a second animal. Scale bar in upper left image is 40 μm. Dark points in scatterplots denote data from cell bodies, lighter points denote data from the neuropil.

## References

Beach, D., & Jacobson, M. (1979). Patterns of cell proliferation in the retina of the clawed frog during development. Journal of Comparative Neurology, 183(3), 603–613.

Brainard, D. H. (1997). The Psychophysics Toolbox. Spatial Vision, 10(4), 433–436.

Chen, T.-W., Wardill, T. J., Sun, Y., Pulver, S. R., Renninger, S. L., Baohan, A., Schreiter, E. R., Kerr, R. A., Orger, M. B., Jayaraman, V., Looger, L. L., Svoboda, K., & Kim, D. S. (2013). Ultrasensitive fluorescent proteins for imaging neuronal activity. Nature, 499(7458), 295–300.

Chung, S.-H., Keating, M., & Bliss, T. (1974). Functional synaptic relations during the development of the retino-tectal projection in amphibians. Proceedings of the Royal Society of London. Series B. Biological Sciences, 187(1089), 449–459.

Cline, H. T., & Kelly, D. (2012). Xenopus as an experimental system for developmental neuroscience: introduction to a special issue. In (Vol. 72, pp. 463–464).

Coupé, P., Munz, M., Manjón, J. V., Ruthazer, E. S., & Collins, D. L. (2012). A CANDLE for a deeper in vivo insight. Medical image analysis, 16(4), 849–864.

Cuntz, H., Forstner, F., Borst, A., & Häusser, M. (2010). One rule to grow them all: a general theory of neuronal branching and its practical application. PLoS computational biology, 6(8), e1000877.

Fedorov, A., Beichel, R., Kalpathy-Cramer, J., Finet, J., Fillion-Robin, J.-C., Pujol, S., Bauer, C., Jennings, D., Fennessy, F., & Sonka, M. (2012). 3D Slicer as an image computing platform for the Quantitative Imaging Network. Magnetic resonance imaging, 30(9), 1323–1341.

Gaze, R., Chung, S., & Keating, M. (1972). Development of the retinotectal projection in Xenopus. Nature New Biology, 236(66), 133–135.

Gaze, R., Keating, M., Ostberg, A., & Chung, S. (1979). The relationship between retinal and tectal growth in larval Xenopus: implications for the development of the retino-tectal projection. Development, 53(1), 103–143.

Gaze, R. M., Keating, M., & Chung, S. (1974). The evolution of the retinotectal map during development in Xenopus. Proceedings of the Royal Society of London. Series B. Biological Sciences, 185(1080), 301–330.

Haas, K., Jensen, K., Sin, W. C., Foa, L., & Cline, H. T. (2002). Targeted electroporation in Xenopus tadpoles in vivo–from single cells to the entire brain. Differentiation, 70(4-5), 148–154.

Haas, K., Sin, W.-C., Javaherian, A., Li, Z., & Cline, H. T. (2001). Single-cell electroporationfor gene transfer in vivo. Neuron, 29(3), 583–591.

Herrgen, L., & Akerman, C. J. (2016). Mapping neurogenesis onset in the optic tectum of Xenopus laevis. Developmental neurobiology, 76(12), 1328–1341.

Hollyfield, J. G. (1971). Differential growth of the neural retina inXenopus laevis larvae. Developmental biology, 24(2), 264–286.

Holt, C. E., & Harris, W. A. (1983). Order in the initial retinotectal map in Xenopus: a new technique for labelling growing nerve fibres. Nature, 301(5896), 150–152.

Kleiner, M., Brainard, D., & Pelli, D. (2007). What’s new in Psychtoolbox-3?

Kutsarova, E., Munz, M., & Ruthazer, E. S. (2017). Rules for Shaping Neural Connections in the Developing Brain. Frontiers in neural circuits, 10, 111–111.

Lazar, G. (1973). The development of the optic tectum in Xenopus laevis: a Golgi study. Journal of anatomy, 116(Pt 3), 347.

Li, V. J., Schohl, A., & Ruthazer, E. S. (2022). Topographic map formation and the effects of NMDA receptor blockade in the developing visual system. Proceedings of the National Academy of Sciences, 119(8), e2107899119.

Liu, Z., Hamodi, A. S., & Pratt, K. G. (2016). Early development and function of the Xenopus tadpole retinotectal circuit. Current Opinion in Neurobiology, 41, 17–23.

McFarlane, S., & Lom, B. (2012). The Xenopus retinal ganglion cell as a model neuron to study the establishment of neuronal connectivity. Developmental Neurobiology, 72(4), 520–536.

McKeown, C. R., & Cline, H. T. (2019). Nutrient restriction causes reversible G2 arrest in Xenopus neural progenitors. Development, 146(20), dev178871.

Molotkov, D., Ferrarese, L., Boissonnet, T., & Asari, H. (2023). Topographic axonal projection at single-cell precision supports local retinotopy in the mouse superior colliculus. Nature communications, 14(1), 7418.

Munz, M., Gobert, D., Schohl, A., Poquerusse, J., Podgorski, K., Spratt, P., & Ruthazer, E. S. (2014). Rapid Hebbian axonal remodeling mediated by visual stimulation. Science, 344(6186), 904–909.

Nieuwkoop, P. D., & Faber, J. (1994). Normal Table of Xenopus Laevis (Daudin): A Systematical and Chronological Survey of the Development from the Fertilized Egg Till the End of Metamorphosis. Garland Pub.

Pachitariu, M., Stringer, C., Schröder, S., Dipoppa, M., Rossi, L. F., Carandini, M., & Harris, K. D. (2016). Suite2p: beyond 10,000 neurons with standard two-photon microscopy. bioRxiv, 061507.

Pelli, D. G. (1997). The VideoToolbox software for visual psychophysics: Transforming numbers into movies. Spatial Vision, 10, 437–442.

Pnevmatikakis, E. A., & Giovannucci, A. (2017). NoRMCorre: An online algorithm for piecewise rigid motion correction of calcium imaging data. Journal of Neuroscience Methods, 291, 83–94.

Roth, G., & Walkowiak, W. (2015). The influence of genome and cell size on brain morphology in amphibians. Cold Spring Harbor perspectives in biology, 7(9), a019075.

Sakaguchi, D. S., & Murphey, R. K. (1985). Map formation in the developing Xenopus retinotectal system: an examination of ganglion cell terminal arborizations. Journal of Neuroscience, 5(12), 3228–3245.

Straznicky, K., & Gaze, R. (1971). The growth of the retina in Xenopus laevis: an autoradiographic study. Development, 26(1), 67–79.

Straznicky, K., & Gaze, R. (1972). The development of the tectum in Xenopus laevis: an autoradiographic study. Development, 28(1), 87–115.

Swain, M. J., & Ballard, D. H. (1991). Color indexing. International journal of computer vision, 7(1), 11–32.

Wu, G.-Y., Malinow, R., & Cline, H. (1996). Maturation of a central glutamatergic synapse. Science, 274(5289), 972–976.

